# Retinoids and EZH2 inhibitors cooperate to orchestrate cytotoxic effects on bladder cancer cells

**DOI:** 10.1101/2023.04.19.537500

**Authors:** Gizem Ozgun, Tutku Yaras, Nick Landman, Gökhan Karakülah, Maarten van Lohuizen, Serif Senturk, Serap Erkek Ozhan

## Abstract

Emerging evidence has highlighted the importance of targeting EZH2 in bladder cancer owing to the highly mutated nature of bladder cancers harboring mutations in chromatin regulatory genes opposing Polycomb-mediated repression. Besides, enhanced expression of EZH2 contributes to pathogenesis. Furthermore, the critical role of the retinoic acid signaling pathway in the development and homeostasis of the urothelium is well established. Here we report that coordinated targeting of EZH2 and the retinoic acid signaling pathway caused cytotoxic effects on bladder cancer cells by inducing a synergistic reduction in proliferative potential that was associated with increased apoptosis and cell cycle arrest in a cooperative and orchestrated manner. Moreover, combined treatment caused the modulation of the expression of genes associated with an anti-oncogenic profile, as reflected by the stimulation of marker genes associated with apoptosis and differentiation. We further portrayed a molecular mechanism whereby EZH2 maintains H3K27me3-mediated repression of certain genes associated with unfolded protein response and some metabolic processes. This work also characterized an apoptotic program centered on the master transcriptional regulators C/EBPβ and CHOP. These findings highlight the importance of co-targeting the EZH2/retinoic acid pathway in bladder cancers and encourage the design of novel treatments employing retinoids coupled with EZH2 inhibitors in bladder carcinoma.

## INTRODUCTION

The substantially proliferative, multifocal, genomically divergent mutations harboring, and metastatic nature of muscle invasive bladder cancer (MIBC) underlies the poor prognosis of the disease ^1^. Owing to the recurrence and frequent resistance to traditional therapies, developing new therapeutic options for bladder cancer patients is required.

Advances in tumor genomic analysis have unveiled dominant mutational signatures of bladder cancers. Bladder cancer has a highly enriched mutational load, with mutations particularly in chromatin regulatory genes encoding proteins such as histone demethylase KDM6A, histone methyltransferases (KMT2C and KMT2D), histone acetyltransferase genes (CREBBP and EP300), and the SWI/SNF chromatin remodelling complex (ARID1A, SMARCA4) ^2–5^. Remarkably, almost all these chromatin modifiers are involved in active chromatin organization, opposing Polycomb mediated repression. The excessive mutational load of chromatin regulatory genes in bladder carcinoma is a hallmark of this disease and a potential weakness that epigenetic drugs could target.

Polycomb group (PcG) proteins function as epigenetic regulators that procure the pursuance of cell-specific transcriptional programs by playing a major role in the implementation of cellular identity and cell fate decisions ^6^. Polycomb repressive complexes (PRCs) are composed of major chromatin modifiers that negatively regulate gene transcription by modulating the global epigenetic state ^7^. The enhancer of zeste homolog 2 (EZH2) methyltransferase is the catalytic subunit of polycomb repressive complex 2 (PRC2), functioning as a transcriptional repressor through trimethylation of lysine 27 of histone 3 (H3K27me3) ^8^. This epigenetic alteration causes chromatin to be densely packed, rendering it hardly accessible to the transcriptional machinery, consequently, eventuates in silencing of genes ^9^. It is established that EZH2 alters the tumor metastatic landscape, affects the tumor microenvironment, and influences cell fate decisions, propounding the highly context-dependent action of EZH2. Notably, EZH2 has been considered as an oncogenic factor in a variety of cancers ^10–12^. Converging lines of investigation have highlighted the role of EZH2 in regulating cell plasticity and encouraging intratumoral heterogeneity, driving the progression of cancer by promoting invasion and metastasis, and ultimately resulting in poor clinical outcomes ^13, 14^. EZH2 orchestrated with other epigenetic modifiers represses genes linked to differentiation and cell cycle arrest, favoring stemness retention ^12, 15, 16^. As observed by previous studies, increased expression of EZH2 contributes to the metastatic cascade of bladder cancers, and selective loss of H3K27me3 function of EZH2 dominates a notable hampering impact on cancer cell migration and tumor metastasis ^17–19^.

Given the proof of EZH2 gain of function becoming a cancer driver, therapeutic strategies have been developed to target EZH2 using specialized chemical inhibitors ^20^. GSK-126, a potent EZH2 inhibitor, has shown to originate loss of genome wide H3K27 methylation, and reactivation of PRC2-repressed genes ^12^. It has been demonstrated that knockdown of EZH2 decreases tumor growth and cell proliferation in a variety of cancers ^21–24^. In addition, inhibiting EZH2 activity causes cancer cells to undergo selective apoptosis, but not normal cells, thereby portraying it as a promising anti-cancer therapeutic target ^25^.

It is known that the application of epigenetic drugs in a combined fashion with other drugs represents a principally rational strategy to sensitize cancer cells to the treatment and to overcome acquired resistance mechanisms ^26^.

Retinoic acids (RAs) are signaling molecules that constitute genetic communication networks having essential functions in embryonic development, organogenesis, organ homeostasis, as well as cell proliferation, differentiation, and death ^27^. Through their driver roles in apoptosis and differentiation, retinoic acid derivatives-retinoids-have gained prominence in research for the development of innovative cancer therapeutics ^28–30^.

Retinoic acid receptor and retinoid X receptor (RAR/RXR) complex, functioning as transcription factors, regulate transcriptional activation of RA target genes via recruitment of certain co-regulators to the RA response elements ^31^. In the absence of ligands for RAR–RXR dimers, recruitment of histone deacetylases (HDACs) leads to chromatin condensation and eventually repression of target genes. Conversely, upon RAR agonist binding, recruited histone acetyltransferase (HAT) complexes bring along derepression of the target genes, in that status, the PRC2 binding is reduced, thus the H3K27me3 is diminished at these regions, and eventually gene expression is activated, emphasizing PRC2’s potential crucial co-repressor role in RA-mediated gene transcription ^32^. Therefore, Polycomb-group proteins function downstream of the RA signaling pathway; inhibiting PRC2 activity could potentially enhance RA action and also overcome the resistance to RA ^33^.

Considering the importance of retinoids in interplaying among several epigenetic processes by altering multiple signaling pathways, we envision that combining retinoids with EZH2 inhibitors could be a promising approach in bladder cancer therapy. In this study, we elucidate the molecular mechanisms linked with the co-treatment of MIBC cell lines with a retinoid acid analogue, fenretinide and an EZH2 inhibitor, GSK-126, as a potential treatment option for bladder cancer. We report that, fenretinide and GSK-126 combined treatment induces a synergistic reduction in proliferative potential correlated with increased apoptosis and modulation of particular genes related to cell proliferation, apoptosis, cell cycle, and urothelial differentiation in MIBC cells.

## RESULTS

### Fenretinide and GSK-126 treatments impair bladder cancer proliferation

Overexpression of EZH2 is implicated in various cancer types and in general is correlated with worse prognosis and higher metastatic stage ^16, 34–36^. Upon comparing the expression of EZH2 in non-muscle invasive bladder cancers (NMIBC) and muscle invasive bladder cancers (MIBC), we detected a significantly increased level of EZH2 expression in muscle invasive cases, both in primary tumors and cell lines (Supplementary Fig. 1). Therefore, collectively, we envisioned that using these MIBC cell lines would be quite informative to test the effect of single and combinatorial EZH2 inhibitor and fenretinide treatments.

We evaluated the effect of fenretinide and a potent EZH2 inhibitor, GSK-126, on cell proliferation in MIBC cell lines using resazurin survival assays. Treatment of the MIBC cell lines with varying doses of fenretinide and GSK-126 for 48 and 72 hours resulted in a dose-dependent decrease in cell proliferation (Fig. 1A, B, Supplementary Fig. 2). Overall, the responses to the fenretinide treatment were similar for 48 and 72 hours, while the response to the GSK-126 treatment required lower doses for 72 hours (Fig. 1, Supplementary Fig. 2).

**Figure 1.**
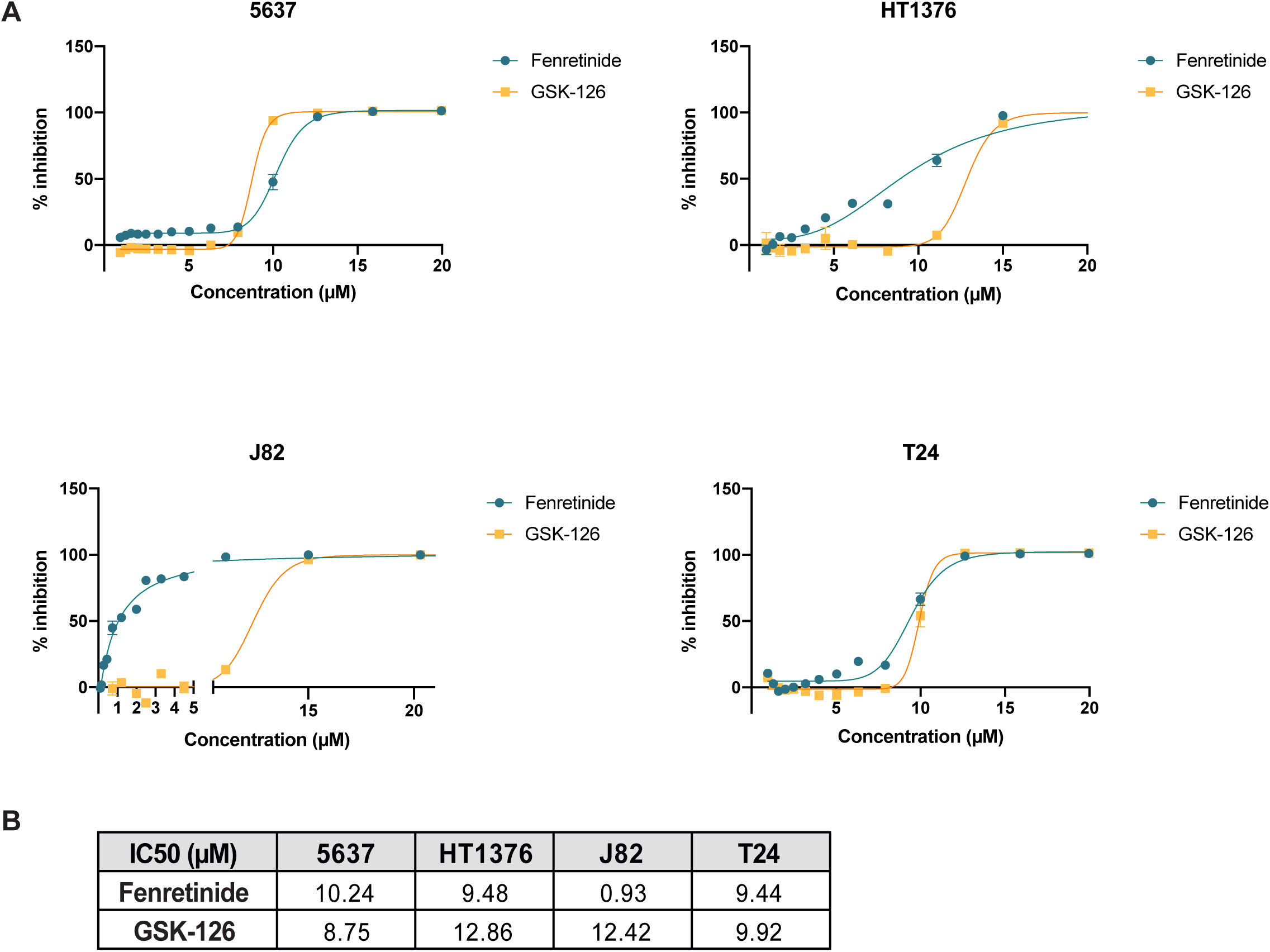
The effects of fenretinide and GSK-126 on MIBC cell viability. a) Dose-response curves of fenretinide and GSK-126 in MIBC cells. Cells were treated for 72 h with increasing concentrations of either fenretinide or GSK-126, and viability was determined in comparison to untreated (DMSO only) control cells. Each point represents the mean and standard deviation of the triplicates. b) IC50 concentrations of MIBC cell lines.

### Fenretinide and GSK-126 combination have a synergic interaction

To investigate retinoid and EZH2 inhibition cooperation, we next designed experiments to study their possible interactions. We tested fenretinide and GSK-126 combination over a range of concentrations to determine whether the drug interactions were additive, synergistic, or antagonistic. Subjecting MIBC cells to increasing concentrations of fenretinide along with GSK-126 showed that this combination suppressed proliferative capacity to a superior extent than either treatment alone. Based on the dose-response results, the synergy scores of this drug combination were calculated as 38.09; 11.46; 4.67; and 57.01 for 5637, HT1376, J82, and T24, respectively, using the Bliss Independence model (Figure 2A,B,C,D). Since a synergy score of >10 is generally accepted as indicating a synergistic interaction ^37–39^, we concluded that fenretinide and GSK-126 showed potential synergistic effects (except for the J82 cell line) (Fig. 2A,B,D). Overall, these results strongly argue for further testing of the combination of fenretinide and GSK-126 against MIBC.

**Figure 2.**
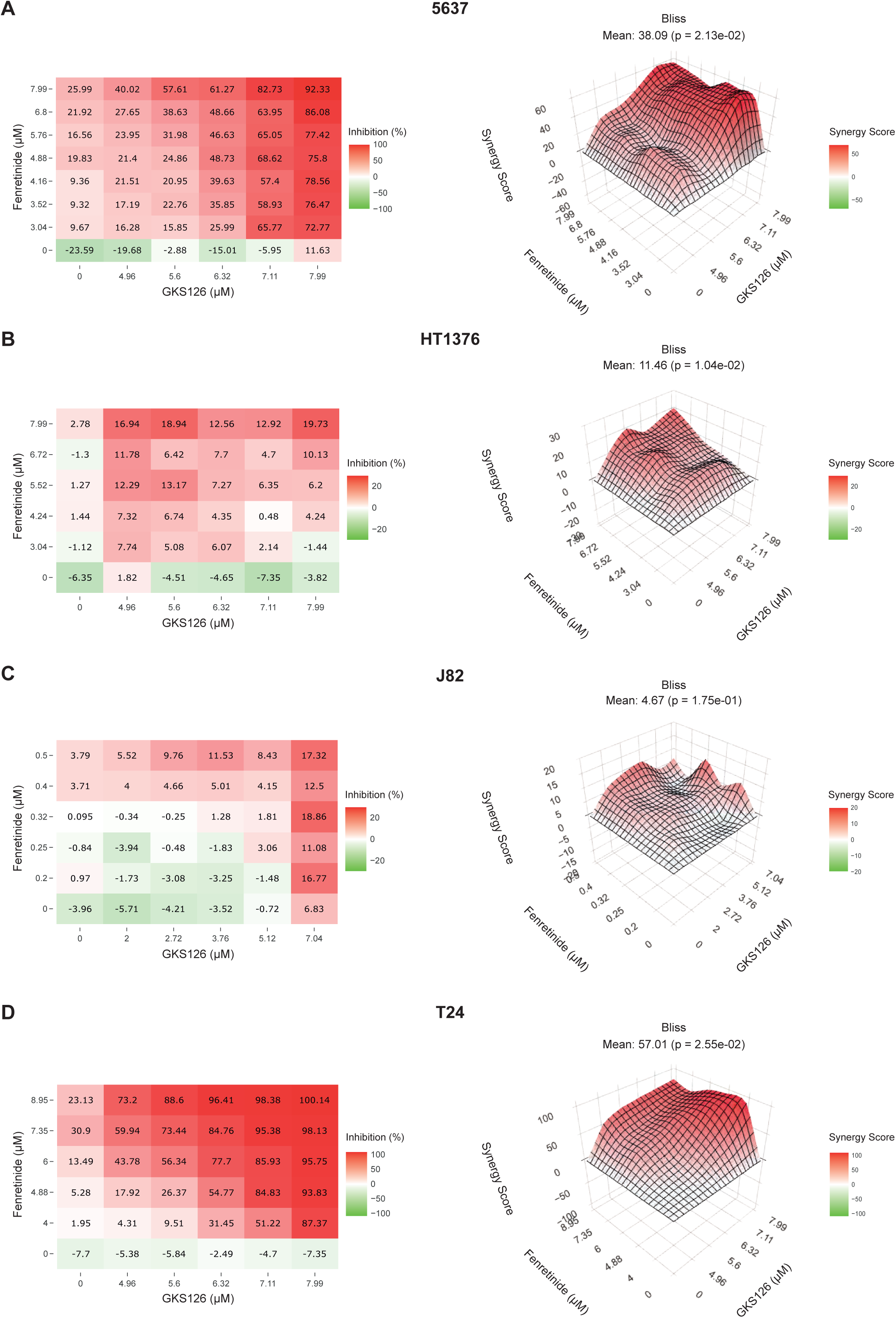
Synergistic effects of fenretinide and GSK-126 on cell viability. Bliss plots showing synergistic effects between fenretinide and GSK-126 in MIBC cells. Bliss Independence synergy scores for the combination of fenretinide and GSK-126 in a 2–8 µM concentration range were indicated. Cells were treated with either single drugs or a drug combination for 72 h. Cell viability (each treatment in triplicate) was assessed by comparison to untreated (DMSO only) control cells. The color of the surface indicates the drug combination effect; green indicates no effect, whereas red indicates maximum effect.

### Fenretinide and GSK-126 combination modulates cell cycle progression, induces apoptosis, inhibits cell migration, and regulates gene expression

Keeping in mind that molecular analysis should be performed at effective doses that do not kill the cells completely, we have chosen 5 µM fenretinide only, 5 µM GSK-126 only, and 5 µM fenretinide and 5 µM GSK-126 combination doses, which correspond to IC25-30 values, to continue our further analysis.

To examine whether the reduction in cell viability was due to apoptotic cell death, we stained the cells with Annexin V-FITC/propidium iodide (PI) after 72 h of treatment with drugs and their combinations. We observed a significant induction of apoptosis with co-treatment with fenretinide and GSK-126, compared with single treatments (Fig. 3A). This finding was also validated by microscopy images, showing the presence of apoptotic cells as revealed by membrane blebbing (Supplementary Fig. 3). Further, the cell cycle distribution of MIBC cells was analyzed after treatments with single agents or their combination. Compared to untreated vehicle control, 72 h treatment of T24 cells with drug combination resulted in a decrease in the S phase, a subtle increase in the G1 fraction, and an increase of the sub-G1 fraction, indicating apoptotic DNA fragmentation consistent with the results of apoptosis (Fig. 3B). The findings are in line with our results generated from cell viability tests. Given the significance of cell migration activity for re-epithelialization, a scratch assay was conducted to test the effects of drugs in combination or alone on cell migration (Fig. 3C). Our results showed that the recovered wound area slightly decreased upon GSK-126 treatment at 24 hours after the scratch. A stronger inhibition of migration was detected for fenretinide treatment at 24 hours after the scratch. Notably, co-treatment reduced migratory capacity in a time-dependent manner, especially at 24 hours after the scratch, compared with either drugs alone (Fig. 3C).

**Figure 3.**
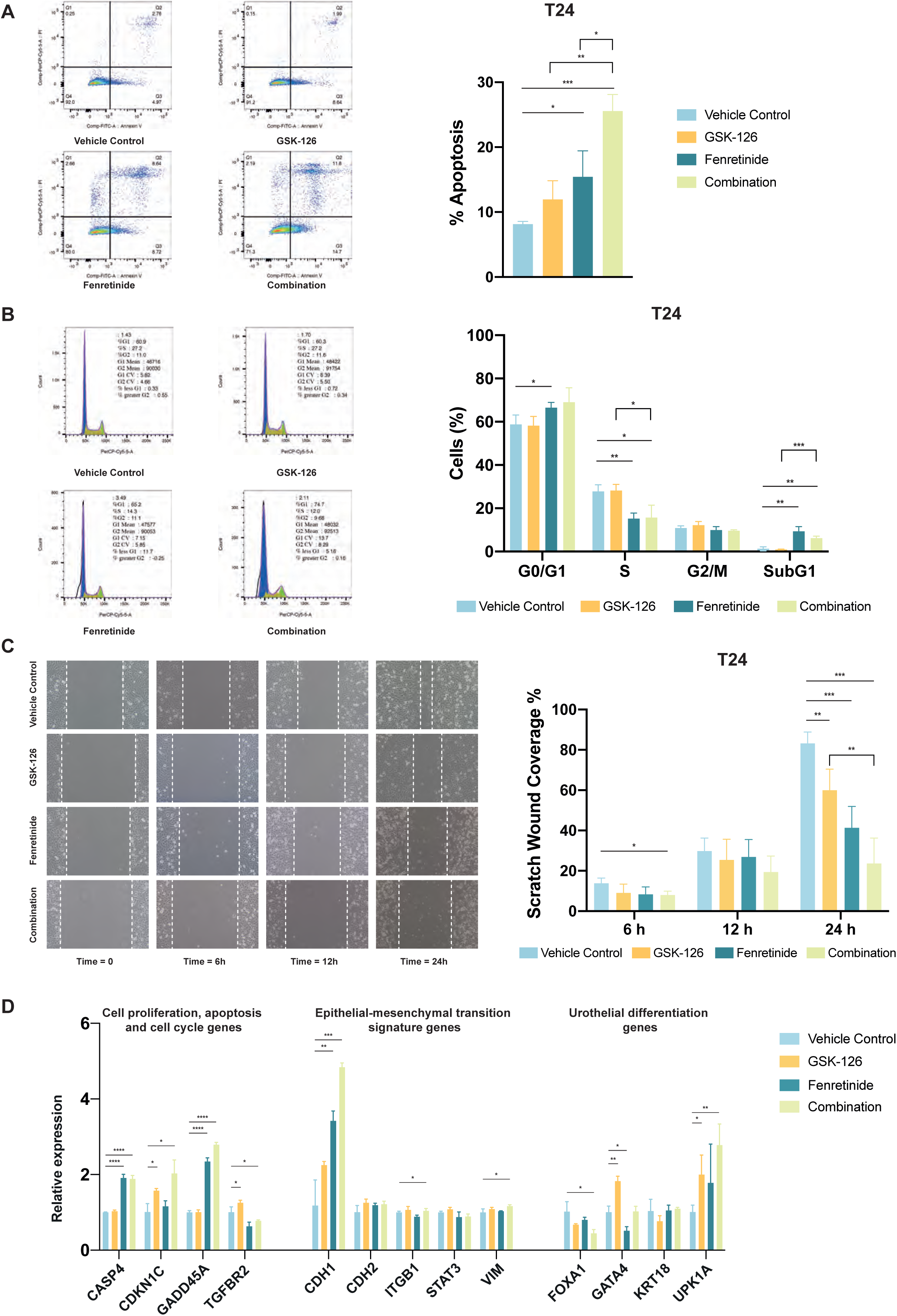
The effects of fenretinide and GSK-126 on cell cycle progression, apoptosis, migration, and gene expression. a) Flow cytometry analyses of apoptosis. T24 cells were treated for 72 h with 5 μM fenretinide, 5 μM GSK-126 or the drug combination. Cell death was assessed by flow cytometry using Annexin-V and PI staining (*p< 0.05, **p < 0.01, ***p < 0.001, two-tailed unpaired t-test, n = 3, Means ± SD are shown). A representative example is on the left. The data summarized is on the right. b) Cell cycle progression as measured by flow cytometry using PI in T24 cells after single or combination treatment for 72 h (representative image from one experiment is on the left). The histogram on the right shows the distribution of cells in G0/G1, S, G2/M, and sub-G1 phases of the cell cycle. Data represent mean ± s.d. (two-tailed unpaired t-test: *p < 0.05, **p < 0.01, ***p < 0.001). c) The effects of drug combination on cell migration. Representative phase-contrast images of the scratch wound migration assay at different time points performed on T24 cell monolayers treated with indicated drugs are on the left. The punctuated white lines indicate the boundaries of the wound. Quantification of the percentage of scratch wound coverage in T24 cells treated with 5 μM fenretinide, 5 μM GSK-126 or the drug combination at different time points. Data represent mean ± s.d. (two-tailed unpaired t-test: *p < 0.05, **p < 0.01, ***p < 0.001). d) The effects of drug combination on gene expression. RT-qPCR analysis of mRNA levels of indicated genes in T24 cells following 72 h treatment with 5 μM fenretinide, 5 μM GSK-126 or the drug combination. Gene specific data were normalized to GAPDH expression and are represented as average relative expression compared to vehicle controls. Error bars specify standard deviation between triplicates. *p< 0.1, **p < 0.01, ***p < 0.001, ****p < 0.0001. p-value was determined by two-tailed, unpaired t-test.

To gain more insights into how the anti-tumor efficacy of fenretinide and GSK-126 in bladder cancer is reflected in gene expression changes, we performed RT-qPCR analysis. We evaluated the changes in gene expression levels in the T24 cell line following treatment with fenretinide, GSK-126, and their combination (Fig. 3D). Confirming our results obtained with the apoptosis assays (Fig. 3A), expression levels of pro-apoptotic genes, CASP4 and GADD45A ^40, 41^ were significantly increased after fenretinide only or fenretinide and GSK-126 combinatorial treatment, suggesting the major role of fenretinide associated with apoptosis. Expression of CDKN1C, implicated in G1 phase cell cycle arrest ^42^, was significantly increased with GSK-126 and combinatorial treatment. Regarding the expression of a subset of EMT marker genes, we observed an effect on the expression level of the epithelial gene CDH1 ^43^ (Fig. 3D). Treatment with the drug combinations resulted in a significant increase in the expression level of CDH1, where maximum changes were observed with the dual treatment of fenretinide and GSK-126, emphasizing the synergistic effect, which might be important for the regulation of this gene. Among the genes related to urothelial differentiation, expression changes for FOXA1 and GATA4 were not uniform; however, expression of UPK1A was significantly increased in a trend similar to that observed for CDH1 (Fig. 3D).

Our findings suggest that fenretinide and GSK-126 cooperate to regulate the expression of the genes critical for apoptosis, urothelial differentiation, and bladder cancer progression, emphasizing the significant anti-oncogenic effects of targeting EZH2 in conjunction with the retinoic acid pathway in MIBC models. Therefore, simultaneous targeting of PRC2 inhibition and the retinoic acid pathway might be a viable strategy for MIBC treatment.

### Fenretinide, GSK-126 alone, and combination treatments result in a distinct set of differentially regulated genes

After observing changes in the expression of key genes involved in apoptosis, cell migration, and cell cycle by RT-qPCR analysis, we decided to investigate the effects on gene regulatory networks associated with fenretinide and GSK-126 treatments at a genome-wide scale by RNA-seq. In our transcriptome analysis, a total of 4308 common protein coding genes were mapped and found to be differentially expressed in T24 cells. Hierarchical clustering analysis of the differentially expressed genes with Log2FC +1 criteria revealed five groups of differentially expressed genes as a result of single and combinatorial treatments compared to control (Fig. 4A). In the light of information acquired from the gene ontology (GO) analysis (p and q values <0.05), the first cluster, combination strongly downregulated (CoSDR), consisted of the genes involved in RNA processing and mitotic spindle organization (Fig. 4B). The second cluster, commonly downregulated after combination or fenretinide-only treatments (CoFeDR), involved the genes regulating chromatin organization, DNA replication, and related processes (Fig. 4B). Genes upregulated after GSK-126 treatment (GskUR, cluster 3) consist of the genes involved in general developmental functions (Figure 4b). The fourth cluster, commonly upregulated after fenretinide only or combination treatments (CoFeUR), was involved in apoptosis and the endoplasmic reticulum (ER) stress response (Fig. 4B). There were only a limited number of genes upregulated as a result of fenretinide-only treatment (FeUR, cluster 5), and those genes were not identified to be involved in a specific pathway. Overall, gene expression changes induced by fenretinide or GSK-126 alone or dual treatments support our phenotypic responses, where apoptotic pathways are triggered while proliferation and cell cycle processes are attenuated.

**Figure 4.**
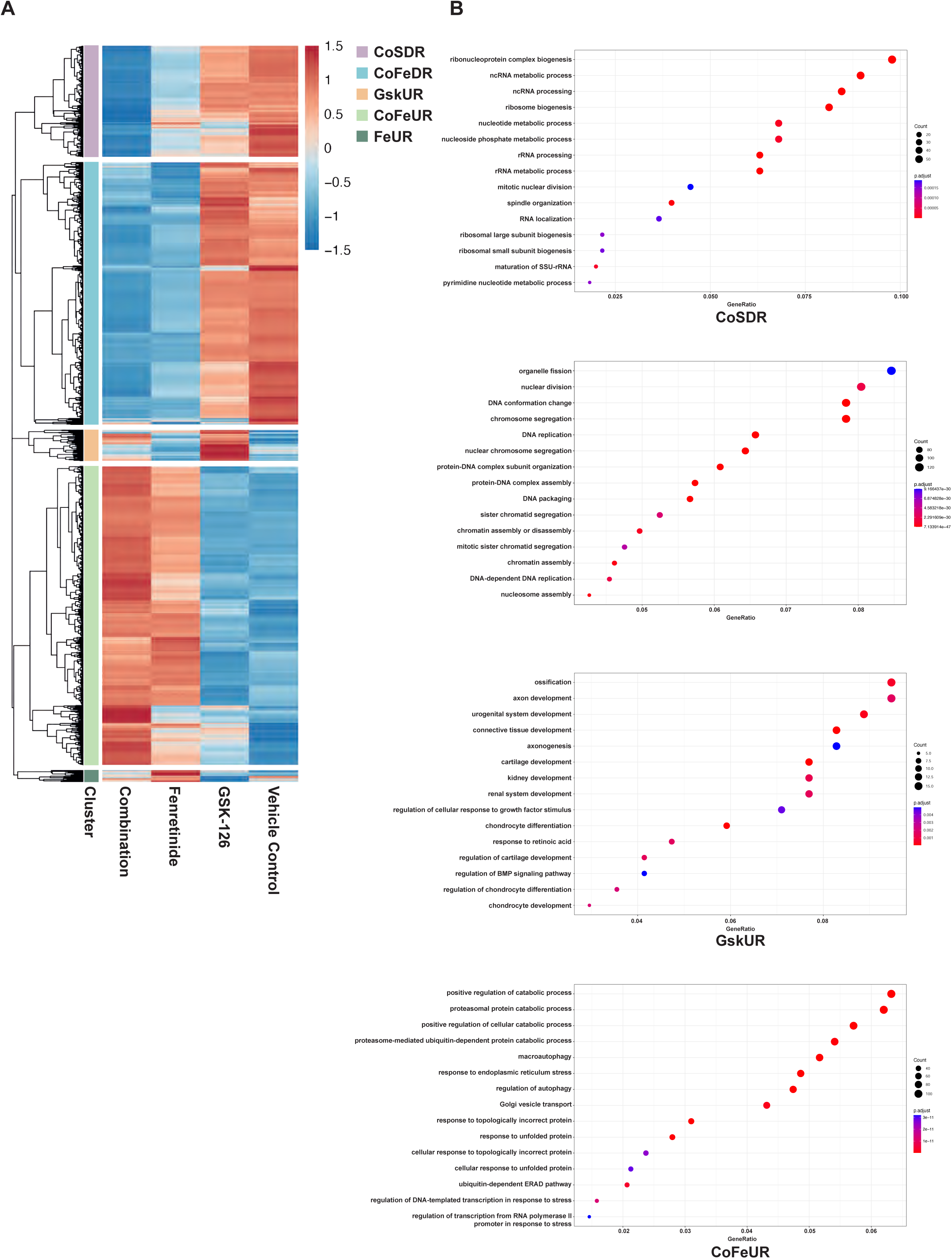
Hierarchical clustering of differentially expressed genes. a) Heat map depicting hierarchical clustering of genes differentially expressed between T24 cells treated with 5 μM fenretinide and 5 μM GSK-126 alone or in combination for 72h versus control using fold change>0.5 cutoff. Red and blue indicate high and low expression of genes, respectively. b) GO enrichment analysis demonstrating the up-regulated biological processes between clusters.

### Association of single and combinatorial treatments with distinct transcription factor activity

To identify the factors potentially regulating distinct classes of differentially expressed genes after fenretinide, GSK-126 only, or combination treatments, we checked the association of differentially expressed genes with transcription factor activities using the information available in the EnrichR database. For the genes belonging to CoSDR and CoFeDR clusters, E2F family transcription factors were significantly enriched, explaining the downregulation of the genes involved in cell cycle-related processes in these clusters (Fig. 5A). Further, generally, expression levels of activator E2F1 and E2F2 transcription factors themselves were strongly downregulated, whereas canonical repressor (E2F3B, E2F5 and E2F6) levels were strongly upregulated after fenretinide only or combination treatment (Supplementary Fig. 4A) ^44^.

**Figure 5.**
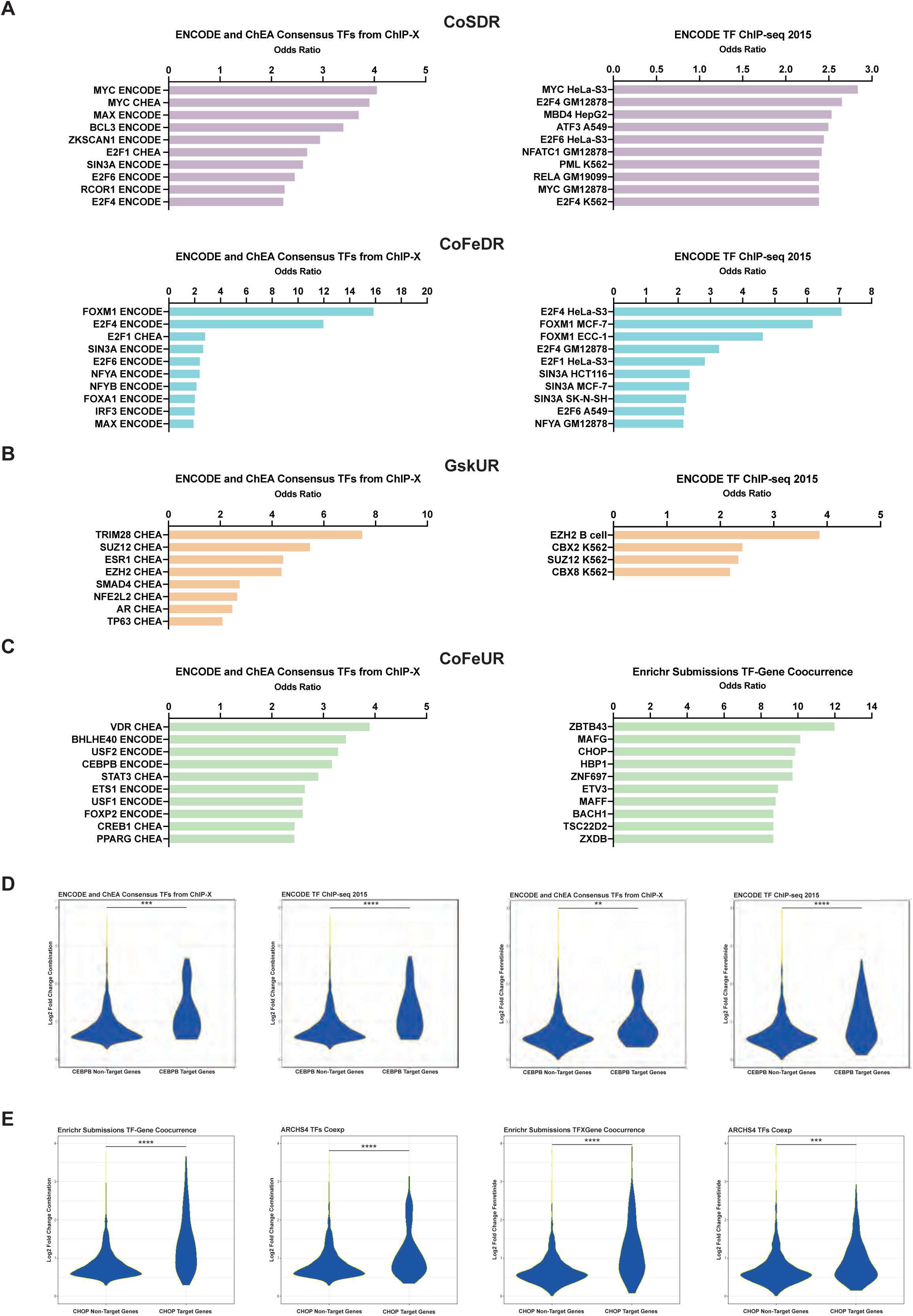
Transcription factor activities of differentially expressed genes. a,b,c) The association of differentially expressed genes with transcription factor activities was determined using the information available in the EnrichR database. DDIT3 was replaced with CHOP for consistency. Adjusted P value < 0.05. d,e) Increased expression of predicted target genes for C/EBPβ and CHOP. The comparison of C/EBPβ and CHOP target and non-target genes which were determined by data obtained from the EnrichR database.

We identified the genes in the GskUR cluster as predominant Polycomb targets (Figure 5b), while the genes belonging to the CoFeUR cluster were enriched for ATF transcription factor family and C/EBP family (Fig. 5C). Importantly, C/EBPβ (CCAAT Enhancer Binding Protein Beta) and DDIT3 (C/EBP homologous protein, CHOP) were among the genes substantially upregulated after fenretinide only and combination treatments. We additionally confirmed the increased level of expression for C/EBPβ and CHOP at protein level (Supplementary Fig. 4B). The roles of these transcription factors are well defined in the ER stress response and apoptosis. For the CoFeUR cluster, enrichment for other transcription factors such as USF1, USF2, NFE2, STAT3, and VDR were not considered to be important as our gene expression data did not show any changes for these factors. We further identified EP300, which is in association with C/EBPβ, as being enriched in CoFeUR cluster genes as well (Supplementary Fig. 4C). Additionally, we discovered that target genes of C/EBPβ and CHOP (DDIT3) were significantly more upregulated in fenretinide only or combination treatments as compared to non-target genes of these two factors (Fig. 5D, E), strengthening the relevance of C/EBPβ and CHOP in the regulation of genes in response to fenretinide and combination treatment.

### GSK-126 treatment in combination with fenretinide enhances the unfolded protein response

Next, we analyzed the CoFeUR cluster more thoroughly by dividing the upregulated genes as combination-enhanced and fenretinide-driven (Fig. 6A). Gene-concept network analysis revealed that combination-enhanced genes were associated with unfolded protein response (UPR), cellular responses to stimuli, and some metabolic processes (Fig. 6B), consisting of the genes, ATF3, DNAJB1, HSPA1A, HSPA1B, and HSPA5. We additionally tested the expression level of some of UPR genes in another MIBC cell line, 5637, showing the second best synergy score for the combination treatment (Supplementary Fig. 5). These results suggest that UPR plays an important role in the synergistic effect of fenretinide and GSK-126. UPR is one major manifestation of ER stress. In this pathway, several branches of signaling result in the activation of CHOP protein. In turn, CHOP triggers the expression of genes involved in apoptosis. It is known that CHOP can form heterodimers with other members of the C/EBP family ^45^. C/EBPβ was previously identified to be involved in apoptosis and ER stress response ^46, 47^.

**Figure 6.**
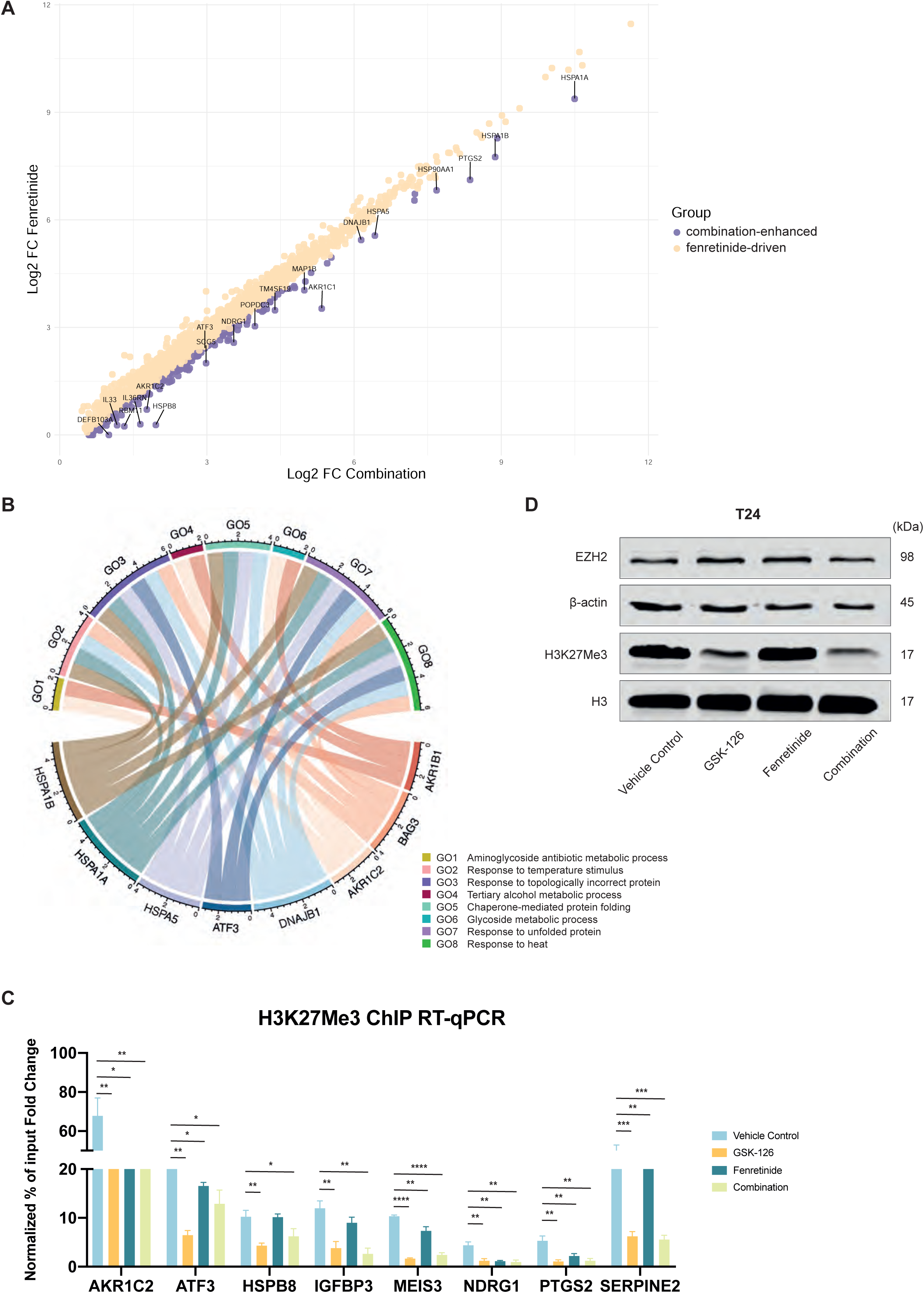
Identification of targets repressed by EZH2. a) Identification of combination-enhanced genes in the CoFeUR cluster. b) Chord diagram of EZH2 targets and their associated GO Biological Terms. c) Differential H3K27me3 levels in T24 cells treated with GSK-126 (5 μM for 72 h) and/or fenretinide (5 μM for 72 h), or DMSO (vehicle). Data were normalized to the negative control GLIS2. d) Western blots performed on T24 cell lines assessing expression of H3K27Me3 and EZH2 after 72 h of treatment with 5 μM fenretinide, 5 μM GSK-126, and the drug combination.

As the genes identified as combination-enhanced in comparison to fenretinide can be viewed as an additional effect brought by GSK-126 over fenretinide, we performed chromatin immunoprecipitation (ChIP) of H3K27me3 for several genes identified as combination-enhanced and determined to be upregulated after GSK-126 only treatment (GskUR) as well. We hypothesized that some of these genes are epigenetically repressed by PRC2 function. Our ChIP-qPCR analysis showed that H3K27me3 levels at AKR1C2, ATF3, HSPB8, IGFBP3, MEIS3, NDRG1, PTGS2, and SERPINE2 genes were significantly reduced by GSK-126 treatment (Fig. 6C). Our data not only identifies these genes as a target of H3K27me3 in MIBC, but also establishes a correlation between the reduction of H3K27me3 with upregulation of gene expression (Fig. 6A). We also validated the character of EZH2 inhibition in line with global reduction of H3K27me3 levels, but not the level of total EZH2 protein. (Fig. 6D).

## DISCUSSION

High-grade bladder tumors harbor considerable risks of recurrence, local invasion, and metastasis. Remarkably, the landscape of therapy regimens for patients with MIBC has not shifted beyond standard management, consisting of radical cystectomy and conventional chemotherapy for nearly 20 years ^12^. Besides, monotherapies largely have limited efficacy and rapidly engender drug resistance. Therefore, providing novel treatment options is urgently needed for MIBC patients.

In this study, we developed a novel approach to target MIBC using the combined action of EZH2 methyltransferase inhibition and the RA signaling pathway as a conceptual framework. We illustrated that the combined effects of EZH2 inhibition and retinoids have anti-oncogenic consequences, and thus this approach might be a viable strategy for MIBC treatment. Notably, we have demonstrated that combined treatment led to an increase in the expression of marker genes associated with apoptosis, cell cycle arrest, and differentiation.

Collectively, the existing literature and our findings highlight the effectiveness of simultaneous targeting of PRC2 and RA signaling in many cancers. Villa and colleagues demonstrated that RA treatment can reverse most of the epigenetic silencing caused by PML-RARα fusion protein, which recruits PRC2 to specific target genes in leukemia patients ^48^. Recently, Benoit and colleagues showed PRC2 inhibition may overcome RA resistance ^49^. RA treatment causes rapid removal of SUZ12, a crucial protein for the assembly of the PRC2, from RA-responsive target genes ^50^. According to research by Ferguson et al. RA signaling and the absence of EZH2 synergistically induce the expression of anti-osteogenic factors ^51^.

Despite progress in the conception of the potent action of retinoids in the treatment or prevention of cancer, there is still a lack of understanding of downstream targets that highlight their biological effects. We demonstrated that fenretinide and combination treatments induce C/EBPβ, CHOP and EP300 expressions thereby triggering anti-oncogenic transcriptional reprogramming resulting in enhanced apoptosis in bladder cancer. In support of all the pre-existing information, our analyses further show that C/EBPβ and CHOP transcription factors conduct their anti-oncogenic functions by stimulating apoptosis and the ER stress response. This is directly compatible with findings from previous studies that prove the role of C/EBPβ /CHOP in apoptosis and the ER stress response ^45–47, 52^. The Duprez group reports that C/EBPβ is a prominent ATRA-dependent target gene implicated in the RA-induced differentiation of acute promyelocytic leukemia cells ^53^. Recently, the correlation between retinoids and C/EBPβ expressions was established ^54, 55^. Our results, highlighting C/EBPβ -mediated apoptosis as a regulatory node for the anti-oncogenic effects of fenretinide, support this conception. Another crucial direction of our results is the positioning of CHOP as one of the putative apoptotic regulators in MIBC. CHOP is widely regarded as a primary controller of ER stress-induced apoptosis. Typically, the UPR and integrated stress response trigger the expression of CHOP. Underpinning our observations, the Gery group has reported CHOP as a novel retinoid-responsive gene during granulocytic differentiation ^56^.

Our hypothesis is C/EBPβ stimulates apoptosis by reassembling accessible chromatin profiles in MIBC cells. Alterations in chromatin architecture conducting apoptotic transcriptional reorientation underlie synergistic interactions. Here, we report that reconfiguration of the apoptotic program is coupled to EZH2. Other groups have reported that disruption of PRC2 activity can enhance apoptosis ^25^, we further nominate C/EBPβ as a unique downstream target of the RA pathway, which may reflect a molecular circuitry supporting apoptotic reprogramming. The output of EZH2-mediated transcriptional rerouting of C/EBPβ/CHOP results in increased apoptosis and may thereby contribute to an improvement in susceptibility to chemotherapy in bladder cancer.

We also attribute the anti-oncogenic effects of this combination to cell cycle arrest. We observed a prominent downregulation in the expression levels of E2F1 and E2F2 family transcription factors with combination therapy, explaining the repression of the genes implicated in cell cycle-related processes. Recently, Lee and colleagues identified E2F1 as a potential EZH2 regulator and a crucial mediator for bladder cancer progression ^57^. We observed an increase for the canonical repressors E2F3B, E2F5 and E2F6 that creates a negative feedback cycle that momentarily switches off the E2F program in G2 ^44^. Consistent with our findings, recently, the Duprez group highlighted the key role of the E2F/EZH2 axis for RA resistance ^58^. We observed an increase in repressors E2F7 and E2F8 with only EZH2 inhibitor treatment. As previously reported, the E2F family can induce the expression of EZH2, which serves as a key downstream mediator of the pRB-E2F pathway ^14^.

Transcriptome analyses and subsequent validation guide us to identify certain genes as targets of H3K27me3. We specify a mechanism by which EZH2 regulates the repression of subsets of genes primarily involved in UPR function. Our results mapping the EZH2 target genes support the notion that these genes were identified in the clusters as mainly functioning in response to topologically incorrect proteins, chaperone-mediated protein folding, and certain metabolic processes. Underpinning our observations, Arbuckle et al demonstrated that EZH2 inhibition promotes the expression of inflammatory cytokines and mediators (IL-6, IL-8, PTGS2) and transcription factors related to the ER stress response (C/EBPβ, FOS, ATF4, DDIT3) ^59^. Recently, Tiffen and colleagues identified the NDRG1 gene as an EZH2 target ^60^. Schrump group also identified IGFBP3, PTGS2, and SERPINE2 as EZH2 targets ^61^.

A number of preclinical and clinical trials proved the superior efficacy and reduced therapeutic recurrence of EZH2 inhibitors, especially in lymphomas but not in solid tumors ^62^. The overall increment in the efficacy of EZH2 inhibitors in hematologic versus solid malignancies is attributed to the existence of particular mutations that cause catalytically hyperactive EZH2, which renders them susceptible to EZH2 inhibitor treatment. Our observation of the strongest combination’s efficacy in T24 cells may be associated with the presence of KDM6A mutations. Cancers harboring loss-of-function mutations in the SWI/SNF complex and KDM6A have been considered to be sensitive to EZH2-based epigenetic therapy ^63^. Ler and colleagues established that KDM6A mutations eventuate in prolonged repression of PRC2-regulated genes in urothelial bladder carcinoma, pointing to the therapeutic benefit of EZH2 inhibition in KDM6A-null or PRC2-enriched urothelial bladder carcinoma ^64^.

Collectively, in this study, we provide additional insights revealing that EZH2 inhibition in MIBC tumors cooperates with retinoids, causing activation of a network of genes providing tumor inhibition. Intriguingly, the network of co-regulatory factors recruited by RA-EZH2 inhibition cooperation has a principal role in the positive regulation of cellular catabolic processes, autophagy, response to ER stress, response to topologically incorrect or unfolded proteins, regulation of DNA−templated transcription in response to stress. The increased co-regulated targets support the notion that this collaboration favors an anti-oncogenic profile. We hypothesize that PRC2 deficiency and retinoic acid pathway stimulation serve as a “second strike” that makes the anti-oncogenic effects of retinoids more penetrant. Therefore, sustaining apoptosis could provide a mechanistic explanation for the cooperative effect of RA gain and PRC2 loss. We speculate that EZH2 may also function as a transcriptional repressor associated with genes controlled by retinoids. Substantially, our findings unveil a completely new aspect of the RA/EZH2 partnership. The fact that retinoic acid-induced transcriptional activity was induced by inhibiting EZH2 underlies the functional significance of this interaction.

Our observations regarding alterations in the transcriptome and epigenome may not reflect the actual clinical scenario. Based on its potent synergistic effects in cell culture, further validation of this novel drug combination in regard to efficacy, tolerability, and pharmacokinetics in animal models is imperative for broadening our results for use in clinical settings.

### Conclusions

Overall, our findings highlight the potential of incorporating the targeting of retinoic acid pathway into PRC2-mediated alterations in epigenetic signatures. Our strategy appears to be a rational approach for improving the therapeutic efficacy in bladder cancers and contributes to targeted treatment paradigms.

## METHODS

### Cell Culture

5637 (DSMZ ACC 35) and HT-1376 (DSMZ ACC 397) human bladder cancer cell lines were purchased from DSMZ. J82 and T24 human bladder cancer cell lines were purchased from ATCC. HT1376, J82, and T24 cells were cultured in DMEM (41965039, Gibco) medium supplemented with 10% fetal bovine serum (10500064, Gibco) and 1% penicillin/streptomycin (15140122, Thermo Fisher), and 5637 cells were cultured in RPMI 1640 (21875, Thermo Fisher) medium supplemented with 10% fetal bovine serum (10500064, Gibco) and 1% penicillin/streptomycin (15140122, Thermo Fisher). The cells were cultured at 37 °C in cell culture dishes in a humidified 5% CO2/air atmosphere. Serial passages were carried out through treatment of sub-confluent monolayers with TrypLE Express Enzyme (12604013, Gibco). All cell lines were confirmed to be mycoplasma-free.

### Drugs and reagents

The EZH2 inhibitor GSK-126 (S7061) was purchased from Selleck Chemicals, USA and Fenretinide (4-Hydroxyphenylretinamide) (390900) was purchased from Merck, Germany. The reagents were dissolved in dimethyl sulfoxide (DMSO) at a concentration of 10 mM and further diluted to different working concentrations.

### Cell viability assays

Cell viability was determined by resazurin survival assays. MIBC cells were seeded on the 384-well plates as 50 μl per well at serial dilution concentrations. Cell number titration is carried out to determine the optimal cell seeding density. After allowing them to attach for 24 h, 10 μl 0.05 mg/ml Resazurin (R7017, Sigma-Aldrich) was added to each well. After 4 hours of incubation at 37 °C, the plate was read on Infinite 200 Pro microplate reader (TECAN) at excitation wavelength 570 nm and emission wavelength 600 nm.

### Drug combination analysis

Prior to cell viability assays, optimal seeding density of cell lines was derived from growth curves. Cells were seeded on 384-well plates in 50 μL of culture medium. After 24 h of incubation, cells were cultured in triplicates in the presence of fenretinide and GSK-126 at serial dilution concentrations of the drugs. DMSO and PAO were used for vehicle control and positive control, respectively. Drug treatments were performed using SPS High Performance D300 Digital Dispenser by HP, Inc. Specialty Printing Systems (1070 NE Circle Blvd Corvallis, OR 97330). After 72 h incubation, 10 μL 0.05 mg/ml of resazurin (R7017, Sigma-Aldrich) was added to each well and incubated at 37 °C for 4 h and plates were read using an Infinite 200 Pro microplate reader (TECAN). Synergy score was calculated using SynergyFinder v2.0 web-based application with default parameters for calculating Bliss Independence.

### Apoptosis assay

Apoptosis was detected in flow cytometry by double staining with Annexin V/propidium iodide (PI). MIBC cells were seeded into 6-well plates. After an overnight incubation, the cells were treated with fenretinide (5 μM), GSK-126 (5 μM),) drug combination, and DMSO as vehicle control. After 72 h of incubation, apoptosis assay was performed according to the instructions of the FITC Annexin V/Dead Cell Apoptosis Kit with FITC Annexin V and PI, for flow cytometry (V13242, Invitrogen). The cells were harvested after the incubation period and washed in cold phosphate-buffered saline (PBS), and then resuspended in 100 μL 1X annexin-binding buffer. After adding 5 μL of Annexin FITC and 1 μL of the 100 μg/mL PI working solution to each cell suspension, the cells were incubated at room temperature for 15 minutes. After the incubation period, the stained cells were analyzed by flow cytometry, measuring the fluorescence emission at 530 nm and >575 nm. Flow cytometer data were analyzed with the FlowJo software.

### Cell cycle analysis

Cells were harvested and resuspended in PBS (1X). Cold ethanol has been added dropwise to each sample to a final concentration of 70% for fixation. The cells were incubated for at least 30 min at 4 °C. Fixed cells are washed in PBS and resuspended in 50 µl of a 100 µg/ml stock of RNase A and then, 200 µl of propidium iodide (stock solution 50 µg/mL) and 0.1% Triton X-100 for staining. The cells were protected from light. The data was acquired on a flow cytometer. Cell cycle phases are analyzed using FlowJo software.

### Scratch assay

The cells were cultured on 6-well plates and incubated overnight to generate a confluent monolayer. The cell monolayer was scratched with a P200 pipette tip. After washing with PBS, the cells were cultured with low serum containing media with or without treatments for different time intervals. Five reference points were randomly selected from a single well, and the percentage of scratch wound closure was analyzed by ImageJ software. Three independent replicate experiments were performed for data representation.

### RNA isolation, cDNA synthesis, and RT-qPCR

Subconfluent cultures of each cell line were treated for 72 h with fenretinide, GSK-126, their combination, or vehicle alone. Total RNA was extracted using the NucleoSpin RNA Midi Kit from Macherey-Nagel according to the manufacturer’s instructions, and 1 μg RNA was transcribed using the Maxima First Strand cDNA Synthesis Kit for RT-qPCR (K1641, Thermo Scientific). Quantitative real-time PCR (qRT-PCR) was performed with FastStart Essential DNA Green Master (06402712001, Roche) with 25 ng of cDNA per reaction (also, 20 µM each primer in a 10 µl reaction volume). Relative gene expression was assessed in three technical replicates of three independent biological samples, and results were normalized to the glyceraldehyde-3-phosphate dehydrogenase (GAPDH) by using the comparative cycles of threshold (Ct) method (ΔΔCt) ^65^. RT-qPCR assays were run on a 7500 fast real-time PCR system (Applied Biosystems). The primer sequences used are listed in Supplementary Table 1.

### RNA-seq data analysis

Raw sequencing files in FastQ files were obtained for four groups, which are “Vehicle Control”, “GSK-126”, “Fenretinide”, and “Combination”. The quality control of raw sequencing data was performed with FASTQC (v0.11.9; https://www.bioinformatics.babraham.ac.uk/projects/fastqc/). Adapter removal and quality trimming steps of sequencing reads were done with Trimmomatic (v0.36) ^66^. The Human reference genome (GRCh38) in FASTA format and related gene annotation in General Transfer Format (GTF) were manually downloaded from the GENCODE (Release 32) project web site (https://www.gencodegenes.org). RNA-seq data of each sample were aligned to the human reference genome with Rusbread v2.0.3package ^67^ of R v4.1.2 statistical computing environment (https://www.r-project.org/) with the following settings: *“align(index={index file}, readfile1={input_1.fastq}, readfile2={input2.fastq} type=“rna”, input_format=“gzFASTQ”, output_format=“BAM”, output_file={output file}, nthreads=numParallelJobs)”*. We utilized SAMtools v1.3.1 ^68^ to sort and index all BAM files created in the alignment step. In order to measure the expression levels of genes, we employed the featureCounts function ^69^ of Rsubread package with the following command: *“featureCounts (files = {infile.bam}, annot.ext = “{infile.gtf}”, isGTFAnnotationFile = T, GTF.featureType = “exon”, GTF.attrType = “gene_id”, useMetaFeatures = T, countMultiMappingReads = T, isPairedEnd = T, nthreads = numParallelJobs)”* and FPKM values of each gene across samples were calculated with “fpkm” function of edgeR package (v3.24.3) ^70^. All genes exhibiting log2(fold change)> 0.5 between groups were considered as differentially expressed. The R statistical computation environment was also utilized for graphical representations of the study. The R-package clusterProfiler (version 3.18.0) ^71^ was used to identify and visualize the enrichment of Gene Ontology (GO) terms in the sets of genes of interest. Graphics associated with RNA-seq data were generated with the R-package ggplot2 (https://ggplot2.tidyverse.org/).

### Western blot analysis

Cellular proteins were extracted using complete RIPA lysis buffer (0.15 M NaCl, 1% NP40, 0.5% DOC, 0.1% SDS, 50 mM Tris (pH 7.5), 2 mM EDTA) with protease inhibitors. Protein lysates were cleared by centrifugation at maximum speed at 4 °C for 15 min. Protein concentrations of whole-cell lysates were measured using the BCA assay. For the preparation of nuclear extracts, histone acid extraction technique that typically results in a massive enrichment of histone proteins was performed (ref: Shechter et al). After washing with PBS, the cell pellet was suspended in a hypotonic lysis buffer containing 10 mM Tris–Cl pH 8.0, 1 mM KCl, 1.5 mM MgCl2 and, 1 mM DTT with a complete protease inhibitor cocktail. After incubating cells for 30 min on the rotator at 4 °C, the intact nuclei were pelleted by spinning in 10,000g for 10 min at 4 °C and suspended in 0.2 M H2SO4. After overnight incubation on the rotator, the supernatant containing histones was precipitated with trichloroacetic acid (TCA). After incubation on ice for 30 min, the histone pellet was washed twice with ice-cold acetone to remove acid from the solution. After drying, the histone pellet was dissolved in ddH2O.

Western blot analysis was performed in accordance with standard protocols. Proteins were denatured by boiling in a 1x Laemmli sample buffer for 5 min. Proteins were separated on Bolt 8–10% Bis-Tris Plus gels and transferred to Bio-Rad nitrocellulose membranes. The membranes were blocked in 5% nonfat dry milk in TBST for 1 hour, and then membranes were incubated overnight at 4 °C with the following primary antibodies: EZH2 (1000x, 5246, Cell Signaling), H3K27Me3 (1000x, 07499, Merck, Millipore), KDM6A (1000x, 33510, Cell Signaling), β-Actin (3700, Cell Signaling) and Histone H3 (39763, Active Motif). After washing with TBST, membranes were incubated with following secondary antibodies: Anti-rabbit IgG (H+L) (5151, CST) and IRDye 680LT Donkey anti-Mouse IgG Secondary Antibody (926-68022, LiCor). Protein bands were visualized using the Odyssey CLX imaging system (LICOR Biosciences).

### Chromatin immunoprecipitation

Cells at 80% confluency were fixed with formaldehyde (1% final concentration) for 10 min, and crosslinking was stopped with 0.125 M glycine. To help to produce chromatin lysates, the cells were incubated with two lysis buffers containing a complete protease inhibitor cocktail (Roche, 11697498001) (Paro Rinse 1: 10 mM Tris pH 8.0, 10 mM EDTA pH 8.0, 0.5 mM EGTA, 0.25% Triton X-100 Paro Rinse 2: 10 mM Tris pH 8.0, 1 mM EDTA, 0.5 mM EGTA, 200 mM NaCl). Then, the cells were lysed in sodium dodecyl sulfate (SDS) lysis buffer (0.1% SDS, 0.1% DOC, 1% Triton X, 1 mM EDTA, 500 mM NaCl, 50 mM HEPES/KOH) containing a complete protease inhibitor cocktail (Roche, 11697498001) and homogenized, followed by incubation on ice for 15 min. The cell lysates were sonicated with Covaris sonicator (S220 High Performance Ultrasonicator) (15 cycles of 140 W Peak Power, 15 Duty Factor, 200 cycles/burst for 60 sec) to generate 200–500 bp long chromatin fragments. Following sonication, the lysates were centrifuged at 16,000 × *g* for 10 min at 4 °C to pellet the debris. In total, 50 μl of the sonicated lysate was stored as input control. RNase A (0.2 mg/ml) was added to clear lysates, and after Proteinase K (200 μg/ml) treatment reversal of crosslink was performed with 1% SDS and 100 mM NaCl. Cross-linking was reversed by heating at 55°C for 2 h and 65 °C for 16-18 h. After the sonication quality was confirmed on agarose gel, DNA product was eluted using Zymo DNA Clean and Concentrator Kit (D4034, Zymo Research) and stored as input DNA. Magnetic beads (11204D, Dynabeads M-280 sheep anti-rabbit IgG, Invitrogen, Thermo Fisher Scientific) were pre-blocked with BSA (1 mg/ml) and tRNA (1 mg/ml). Chromatin sample was pre-cleared with tRNA pre-blocked magnetic beads. H3K27me3 antibody (9733, Cell Signaling Technology) was used for immunoprecipitation of chromatin and incubated overnight at 4°C with overhead shaking. After adding corresponding pre-blocked magnetic beads to the AB-chromatin complex, the complex incubated for 3 h at 4°C with overhead shaking. Followed by washing lysis and high salt buffers (0.5% DOC, 0.5 % NP-40, 0.25 M LiCl, 1 mM EDTA, 10 mM Tris pH 8.0), the immunoprecipitated DNA was eluted off the magnetic beads with elution buffer (1 M NaHCO3, 1% SDS) and reverse cross-linked by heating at 55°C for 2 h and 65 °C for 16-18 h. The immunoprecipitated DNA was eluted using a ChIP DNA Clean and Concentrator Kit (D5205, Zymo Research). H3K27me3 content at the promoters of specified genes were assessed by RT-qPCR using primers designed from the Cistrome Data Browser resource ^72^. ChIP primer sequences for different genes are enlisted in Supplementary Table 2. The enrichment of immunoprecipitated DNA with anti-H3K27me3 antibody was calculated using the following method: The average Ct of triplicates was calculated. Adjusted Ct value for input sample was calculated as (Ct_10% Input_-log (10,2)). ΔCt value of input sample was calculated (Ct_Adjusted Input_ -Ct_ChIP Input_), the percent input was calculated as (100∗2^(ΔCt)).

### Quantification, statistical analyses, and reproducibility

All data sets except RNA-seq data were analyzed with GraphPad Prism (version 7.0). All assays were performed in triplicate. Statistical differences between groups were calculated by applying analysis of variance (ANOVA) or a Student’s *t* test, as indicated. *p<0.05, **p<0.01, ***p<0.001, ****p<0.0001.

## ACKNOWLEDGEMENTS

This work was funded by The Scientific and Technological Research Council of Turkey (TÜBİTAK), and the EMBO Installation Grant (No:4148). The international collaboration was supported by EMBO Scientific Exchange Grant (No:8981).

## Contributions

GO, SS, and SEO developed the concept of this study. GO performed all the experiments. GO and TY analyzed and visualized the data. GO and SEO wrote the manuscript. GO, SEO, SS, GK, and MvL contributed to the study design and data interpretation. NL contributed to the method optimization and data acquisition for drug combination experiments. GO, TY, and GK performed bioinformatical analyses. SEO supervised this work and acquired funding. All authors have read and agreed to the published version of the manuscript.

## REFERENCES

1. Sanli O, Dobruch J, Knowles MA, et al. Bladder cancer. Nat Rev Dis Prim. 2017;3(13):17022. doi:10.1038/nrdp.2017.22

2. Robertson AG, Kim J, Al-Ahmadie H, et al. Comprehensive Molecular Characterization of Muscle-Invasive Bladder Cancer. Cell. 2017;171(3):540–556.e25. doi:10.1016/j.cell.2017.09.007

3. The Cancer Genome Atlas Research Network. Comprehensive molecular characterization of urothelial bladder carcinoma. Nature. 2014;507(7492):315-322. doi:10.1038/nature12965

4. Kamoun A, de Reyniès A, Allory Y, et al. A Consensus Molecular Classification of Muscle-invasive Bladder Cancer. Eur Urol. 2020;77(4):420–433. doi:https://doi.org/10.1016/j.eururo.2019.09.006

5. Taber A, Christensen E, Lamy P, et al. Molecular correlates of cisplatin-based chemotherapy response in muscle invasive bladder cancer by integrated multi-omics analysis. Nat Commun. 2020;11(1):4858. doi:10.1038/s41467-020-18640-0

6. Lavarone E, Barbieri CM, Pasini D. Dissecting the role of H3K27 acetylation and methylation in PRC2 mediated control of cellular identity. Nat Commun. 2019;10(1):1679. doi:10.1038/s41467-019-09624-w

7. Piunti A, Shilatifard A. The roles of Polycomb repressive complexes in mammalian development and cancer. Nat Rev Mol Cell Biol. 2021;22(5):326–345. doi:10.1038/s41580-021-00341-1

8. Cha TL, Zhou BP, Xia W, et al. Molecular biology: Akt-mediated phosphorylation of EZH2 suppresses methylation of lysine 27 in histone H3. Science (80-). 2005;310(5746):306–310. doi:10.1126/science.1118947

9. Simon JA, Lange CA. Roles of the EZH2 histone methyltransferase in cancer epigenetics. Mutat Res Mol Mech Mutagen. 2008;647(1):21–29. doi:https://doi.org/10.1016/j.mrfmmm.2008.07.010

10. Ren G, Baritaki S, Marathe H, et al. Polycomb protein EZH2 regulates tumor invasion via the transcriptional repression of the metastasis suppressor RKIP in breast and prostate cancer. Cancer Res. 2012;72(12):3091–3104. doi:10.1158/0008-5472.CAN-11-3546

11. Zhao L, Yu Y, Wu J, et al. Role of EZH2 in oral squamous cell carcinoma carcinogenesis. Gene. 2014;537(2):197–202. doi:https://doi.org/10.1016/j.gene.2014.01.006

12. Martínez-Fernández M, Rubio C, Segovia C, López-Calderón FF, Dueñas M, Paramio JM. EZH2 in Bladder Cancer, a Promising Therapeutic Target. Int J Mol Sci. 2015;16(11):27107–27132. doi:10.3390/ijms161126000

13. Bachmann IM, Halvorsen OJ, Collett K, et al. EZH2 expression is associated with high proliferation rate and aggressive tumor subgroups in cutaneous melanoma and cancers of the endometrium, prostate, and breast. J Clin Oncol. 2006;24(2):268–273. doi:10.1200/JCO.2005.01.5180

14. Bracken AP, Pasini D, Capra M, Prosperini E, Colli E, Helin K. EZH2 is downstream of the pRB-E2F pathway, essential for proliferation and amplified in cancer. EMBO J. 2003;22(20):5323–5335. doi:10.1093/emboj/cdg542

15. Ezhkova E, Pasolli HA, Parker JS, et al. Ezh2 orchestrates gene expression for the stepwise differentiation of tissue-specific stem cells. Cell. 2009;136(6):1122–1135. doi:10.1016/j.cell.2008.12.043

16. Kim KH, Roberts CWM. Targeting EZH2 in cancer. Nat Med. 2016;22(2):128–134. doi:10.1038/nm.4036

17. Chen Z, Du Y, Liu X, et al. EZH2 inhibition suppresses bladder cancer cell growth and metastasis via the JAK2/STAT3 signaling pathway. Oncol Lett. 2019;18(1):907–915. doi:10.3892/ol.2019.10359

18. Zhou X, Liu N, Zhang J, et al. Increased expression of EZH2 indicates aggressive potential of urothelial carcinoma of the bladder in a Chinese population. Sci Rep. 2018;8(1):17792. doi:10.1038/s41598-018-36164-y

19. Liu D, Li Y, Luo G, et al. LncRNA SPRY4-IT1 sponges miR-101-3p to promote proliferation and metastasis of bladder cancer cells through up-regulating EZH2. Cancer Lett. 2017;388:281–291. doi:10.1016/j.canlet.2016.12.005

20. Pfister SX, Ashworth A. Marked for death: targeting epigenetic changes in cancer. Nat Rev Drug Discov. 2017;16(4):241–263. doi:10.1038/nrd.2016.256

21. McCabe MT, Ott HM, Ganji G, et al. EZH2 inhibition as a therapeutic strategy for lymphoma with EZH2-activating mutations. Nature. 2012;492(7427):108–112. doi:10.1038/nature11606

22. Ihira K, Dong P, Xiong Y, et al. EZH2 inhibition suppresses endometrial cancer progression via miR-361/Twist axis. Oncotarget. 2017;8(8):13509–13520. doi:10.18632/oncotarget.14586

23. Bownes L V, Williams AP, Marayati R, et al. EZH2 inhibition decreases neuroblastoma proliferation and in vivo tumor growth. PLoS One. 2021;16(3):e0246244. doi:10.1371/journal.pone.0246244

24. Lv Y-F, Yan G-N, Meng G, Zhang X, Guo Q-N. Enhancer of zeste homolog 2 silencing inhibits tumor growth and lung metastasis in osteosarcoma. Sci Rep. 2015;5(1):12999. doi:10.1038/srep12999

25. Tan J, Yang X, Zhuang L, et al. Pharmacologic disruption of Polycomb-repressive complex 2-mediated gene repression selectively induces apoptosis in cancer cells. Genes Dev. 2007;21(9):1050–1063. doi:10.1101/gad.1524107

26. Morel D, Jeffery D, Aspeslagh S, Almouzni G, Postel-Vinay S. Combining epigenetic drugs with other therapies for solid tumours — past lessons and future promise. Nat Rev Clin Oncol. 2020;17(2):91–107. doi:10.1038/s41571-019-0267-4

27. Gudas LJ, Wagner JA. Retinoids regulate stem cell differentiation. J Cell Physiol. 2011;226(2):322–330. doi:10.1002/jcp.22417

28. Altucci L, Gronemeyer H. The promise of retinoids to fight against cancer. Nat Rev Cancer. 2001;1(3):181–193. doi:10.1038/35106036

29. Chen S, Hu Q, Tao X, et al. Retinoids in cancer chemoprevention and therapy: Meta-analysis of randomized controlled trials. Front Genet. 2022;13:1–14. https://www.frontiersin.org/articles/10.3389/fgene.2022.1065320.

30. Clifford JL, Sabichi AL, Zou C, et al. Effects of novel phenylretinamides on cell growth and apoptosis in bladder cancer. Cancer Epidemiol biomarkers Prev a Publ Am Assoc Cancer Res cosponsored by Am Soc Prev Oncol. 2001;10(4):391–395.

31. le Maire A, Teyssier C, Balaguer P, Bourguet W, Germain P. Regulation of RXR-RAR Heterodimers by RXR- and RAR-Specific Ligands and Their Combinations. Cells. 2019;8(11). doi:10.3390/cells8111392

32. Gillespie RF, Gudas LJ. Retinoid regulated association of transcriptional co-regulators and the polycomb group protein SUZ12 with the retinoic acid response elements of Hoxa1, RARbeta(2), and Cyp26A1 in F9 embryonal carcinoma cells. J Mol Biol. 2007;372(2):298–316. doi:10.1016/j.jmb.2007.06.079

33. Tang X-H, Gudas LJ. Retinoids, retinoic acid receptors, and cancer. Annu Rev Pathol. 2011;6:345–364. doi:10.1146/annurev-pathol-011110-130303

34. Wang H, Albadine R, Magheli A, et al. Increased EZH2 protein expression is associated with invasive urothelial carcinoma of the bladder. Urol Oncol. 2012;30(4):428–433. doi:10.1016/j.urolonc.2010.09.005

35. Pawlyn C, Bright MD, Buros AF, et al. Overexpression of EZH2 in multiple myeloma is associated with poor prognosis and dysregulation of cell cycle control. Blood Cancer J. 2017;7(3):e549–e549. doi:10.1038/bcj.2017.27

36. Kleer CG, Cao Q, Varambally S, et al. EZH2 is a marker of aggressive breast cancer and promotes neoplastic transformation of breast epithelial cells. Proc Natl Acad Sci U S A. 2003;100(20):11606–11611. doi:10.1073/pnas.1933744100

37. Yadav B, Wennerberg K, Aittokallio T, Tang J. Searching for Drug Synergy in Complex Dose-Response Landscapes Using an Interaction Potency Model. Comput Struct Biotechnol J. 2015;13:504–513. doi:10.1016/j.csbj.2015.09.001

38. Zheng S, Wang W, Aldahdooh J, et al. SynergyFinder Plus: Toward Better Interpretation and Annotation of Drug Combination Screening Datasets. Genomics Proteomics Bioinformatics. 2022;20(3):587–596. doi:https://doi.org/10.1016/j.gpb.2022.01.004

39. Lederer S, Dijkstra TMH, Heskes T. Additive Dose Response Models: Defining Synergy. Front Pharmacol. 2019;10:1–15. https://www.frontiersin.org/articles/10.3389/fphar.2019.01384.

40. Hitomi J, Katayama T, Eguchi Y, et al. Involvement of caspase-4 in endoplasmic reticulum stress-induced apoptosis and Aβ-induced cell death . J Cell Biol. 2004;165(3):347–356. doi:10.1083/jcb.200310015

41. Kleinsimon S, Longmuss E, Rolff J, et al. GADD45A and CDKN1A are involved in apoptosis and cell cycle modulatory effects of viscumTT with further inactivation of the STAT3 pathway. Sci Rep. 2018;8(1):5750. doi:10.1038/s41598-018-24075-x

42. Borges KS, Arboleda VA, Vilain E. Mutations in the PCNA-binding site of CDKN1C inhibit cell proliferation by impairing the entry into S phase. Cell Div. 2015;10:2. doi:10.1186/s13008-015-0008-8

43. van Roy F. Beyond E-cadherin: roles of other cadherin superfamily members in cancer. Nat Rev Cancer. 2014;14(2):121–134. doi:10.1038/nrc3647

44. Kent LN, Leone G. The broken cycle: E2F dysfunction in cancer. Nat Rev Cancer. 2019;19(6):326–338. doi:10.1038/s41568-019-0143-7

45. Hu H, Tian M, Ding C, Yu S. The C/EBP Homologous Protein (CHOP) Transcription Factor Functions in Endoplasmic Reticulum Stress-Induced Apoptosis and Microbial Infection. Front Immunol. 2018;9:3083. doi:10.3389/fimmu.2018.03083

46. Zahid MDK, Rogowski M, Ponce C, Choudhury M, Moustaid-Moussa N, Rahman SM. CCAAT/enhancer-binding protein beta (C/EBPβ) knockdown reduces inflammation, ER stress, and apoptosis, and promotes autophagy in oxLDL-treated RAW264.7 macrophage cells. Mol Cell Biochem. 2020;463(1-2):211–223. doi:10.1007/s11010-019-03642-4

47. Yang Y, Liu L, Naik I, Braunstein Z, Zhong J, Ren B. Transcription Factor C/EBP Homologous Protein in Health and Diseases. Front Immunol. 2017;8:1612. doi:10.3389/fimmu.2017.01612

48. Villa R, Pasini D, Gutierrez A, et al. Role of the Polycomb Repressive Complex 2 in Acute Promyelocytic Leukemia. Cancer Cell. 2007;11(6):513–525. doi:https://doi.org/10.1016/j.ccr.2007.04.009

49. Benoit YD, Laursen KB, Witherspoon MS, Lipkin SM, Gudas LJ. Inhibition of PRC2 histone methyltransferase activity increases TRAIL-mediated apoptosis sensitivity in human colon cancer cells. J Cell Physiol. 2013;228(4):764–772. doi:10.1002/jcp.24224

50. Amat R, Gudas LJ. RARγ is required for correct deposition and removal of Suz12 and H2A.Z in embryonic stem cells. J Cell Physiol. 2011;226(2):293–298. doi:10.1002/jcp.22420

51. Ferguson JW, Devarajan M, Atit RP. Stage-specific roles of Ezh2 and Retinoic acid signaling ensure calvarial bone lineage commitment. Dev Biol. 2018;443(2):173–187. doi:10.1016/j.ydbio.2018.09.014

52. Allagnat F, Fukaya M, Nogueira TC, et al. C/EBP homologous protein contributes to cytokine-induced pro-inflammatory responses and apoptosis in β-cells. Cell Death Differ. 2012;19(11):1836–1846. doi:10.1038/cdd.2012.67

53. Duprez E, Wagner K, Koch H, Tenen DG. C/EBPbeta: a major PML-RARA-responsive gene in retinoic acid-induced differentiation of APL cells. EMBO J. 2003;22(21):5806–5816. doi:10.1093/emboj/cdg556

54. Kutty RK, Samuel W, Jaworski C, et al. MicroRNA expression in human retinal pigment epithelial (ARPE-19) cells: increased expression of microRNA-9 by N-(4-hydroxyphenyl)retinamide. Mol Vis. 2010;16:1475–1486.

55. Marchwicka A, Marcinkowska E. Regulation of Expression of CEBP Genes by Variably Expressed Vitamin D Receptor and Retinoic Acid Receptor α in Human Acute Myeloid Leukemia Cell Lines. Int J Mol Sci. 2018;19(7). doi:10.3390/ijms19071918

56. Gery S, Park DJ, Vuong PT, Chih DY, Lemp N, Koeffler HP. Retinoic acid regulates C/EBP homologous protein expression (CHOP), which negatively regulates myeloid target genes. Blood. 2004;104(13):3911–3917. doi:10.1182/blood-2003-10-3688

57. Lee S-R, Roh Y-G, Kim S-K, et al. Activation of EZH2 and SUZ12 Regulated by E2F1 Predicts the Disease Progression and Aggressive Characteristics of Bladder Cancer. Clin cancer Res an Off J Am Assoc Cancer Res. 2015;21(23):5391–5403. doi:10.1158/1078-0432.CCR-14-2680

58. Poplineau M, Platet N, Mazuel A, et al. Noncanonical EZH2 drives retinoic acid resistance of variant acute promyelocytic leukemias. Blood. 2022;140(22):2358–2370. doi:10.1182/blood.2022015668

59. Arbuckle JH, Gardina PJ, Gordon DN, et al. Inhibitors of the Histone Methyltransferases EZH2/1 Induce a Potent Antiviral State and Suppress Infection by Diverse Viral Pathogens. MBio. 2017;8(4):e01141–17. doi:10.1128/mBio.01141-17

60. Tiffen J, Gallagher SJ, Filipp F, et al. EZH2 Cooperates with DNA Methylation to Downregulate Key Tumor Suppressors and IFN Gene Signatures in Melanoma. J Invest Dermatol. 2020;140(12):2442–2454.e5. doi:https://doi.org/10.1016/j.jid.2020.02.042

61. Kemp CD, Rao M, Xi S, et al. Polycomb repressor complex-2 is a novel target for mesothelioma therapy. Clin cancer Res an Off J Am Assoc Cancer Res. 2012;18(1):77–90. doi:10.1158/1078-0432.CCR-11-0962

62. Gulati N, Béguelin W, Giulino-Roth L. Enhancer of zeste homolog 2 (EZH2) inhibitors. Leuk Lymphoma. 2018;59(7):1574–1585. doi:10.1080/10428194.2018.1430795

63. Ramakrishnan S, Granger V, Rak M, et al. Inhibition of EZH2 induces NK cell-mediated differentiation and death in muscle-invasive bladder cancer. Cell Death Differ. 2019;26(10):2100–2114. doi:10.1038/s41418-019-0278-9

64. Ler LD, Ghosh S, Chai X, et al. Loss of tumor suppressor KDM6A amplifies PRC2-regulated transcriptional repression in bladder cancer and can be targeted through inhibition of EZH2. Sci Transl Med. 2017;9(378):eaai8312. doi:10.1126/scitranslmed.aai8312

65. Livak KJ, Schmittgen TD. Analysis of Relative Gene Expression Data Using Real-Time Quantitative PCR and the 2−ΔΔCT Method. Methods. 2001;25(4):402–408. doi:https://doi.org/10.1006/meth.2001.1262

66. Bolger AM, Lohse M, Usadel B. Trimmomatic: a flexible trimmer for Illumina sequence data. Bioinformatics. 2014;30(15):2114–2120. doi:10.1093/bioinformatics/btu170

67. Liao Y, Smyth GK, Shi W. The R package Rsubread is easier, faster, cheaper and better for alignment and quantification of RNA sequencing reads. Nucleic Acids Res. 2019;47(8):e47. doi:10.1093/nar/gkz114

68. Li H, Handsaker B, Wysoker A, et al. The Sequence Alignment/Map format and SAMtools. Bioinformatics. 2009;25(16):2078–2079. doi:10.1093/bioinformatics/btp352

69. Liao Y, Smyth GK, Shi W. featureCounts: an efficient general purpose program for assigning sequence reads to genomic features. Bioinformatics. 2014;30(7):923–930. doi:10.1093/bioinformatics/btt656

70. Robinson MD, McCarthy DJ, Smyth GK. edgeR: a Bioconductor package for differential expression analysis of digital gene expression data. Bioinformatics. 2010;26(1):139–140. doi:10.1093/bioinformatics/btp616

71. Yu G, Wang L-G, Han Y, He Q-Y. clusterProfiler: an R package for comparing biological themes among gene clusters. OMICS. 2012;16(5):284–287. doi:10.1089/omi.2011.0118

72. Liu T, Ortiz JA, Taing L, et al. Cistrome: an integrative platform for transcriptional regulation studies. Genome Biol. 2011;12(8):R83. doi:10.1186/gb-2011-12-8-r83

