## Supplementary Information for "Retinoids and EZH2 inhibitors cooperate to orchestrate cytotoxic effects on bladder cancer cells"

Supplementary Figure 1. Correlation between EZH2 mRNA levels and bladder cancer progression

Supplementary Figure 2. The effects of fenretinide and GSK-126 on MIBC cell viability
.
Supplementary Figure 3. Apoptosis induced by fenretinide and GSK-126 co-treatment

Supplementary Figure 4. The association of fenretinide only and combination treatments with cell cycle-related processes, CEBPB and CHOP protein levels, and transcription factor activities

Supplementary Figure 5. The effects of combination treatment on unfolded protein response

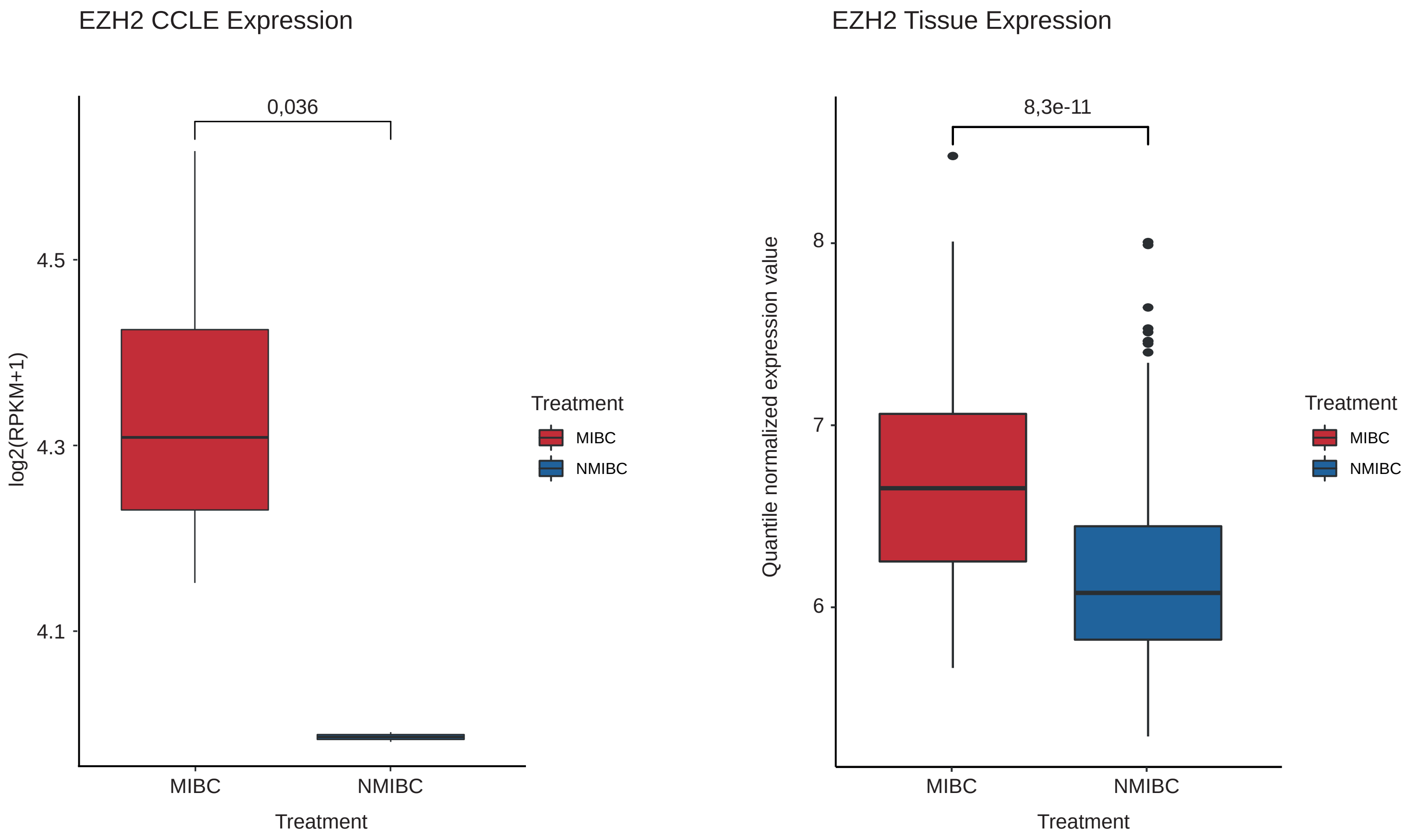


**Supplementary Figure 1. Correlation between EZH2 mRNA levels and bladder cancer progression**
Left: The comparison of EZH2 levels in NMIBC and MIBC cells. Data of EZH2 mRNA expression levels in MIBC cell lines (5637, HT1376, J82 and T24) and NMIBC cell lines (RT4, RT112) were obtained from publicly available Cancer Cell Line Encyclopedia (CCLE) database. Right: The comparison of the EZH2 signature of microarray data of NMIBC and MIBC tissue samples (GEO: GSE32894). P values of t-test are indicated.

A

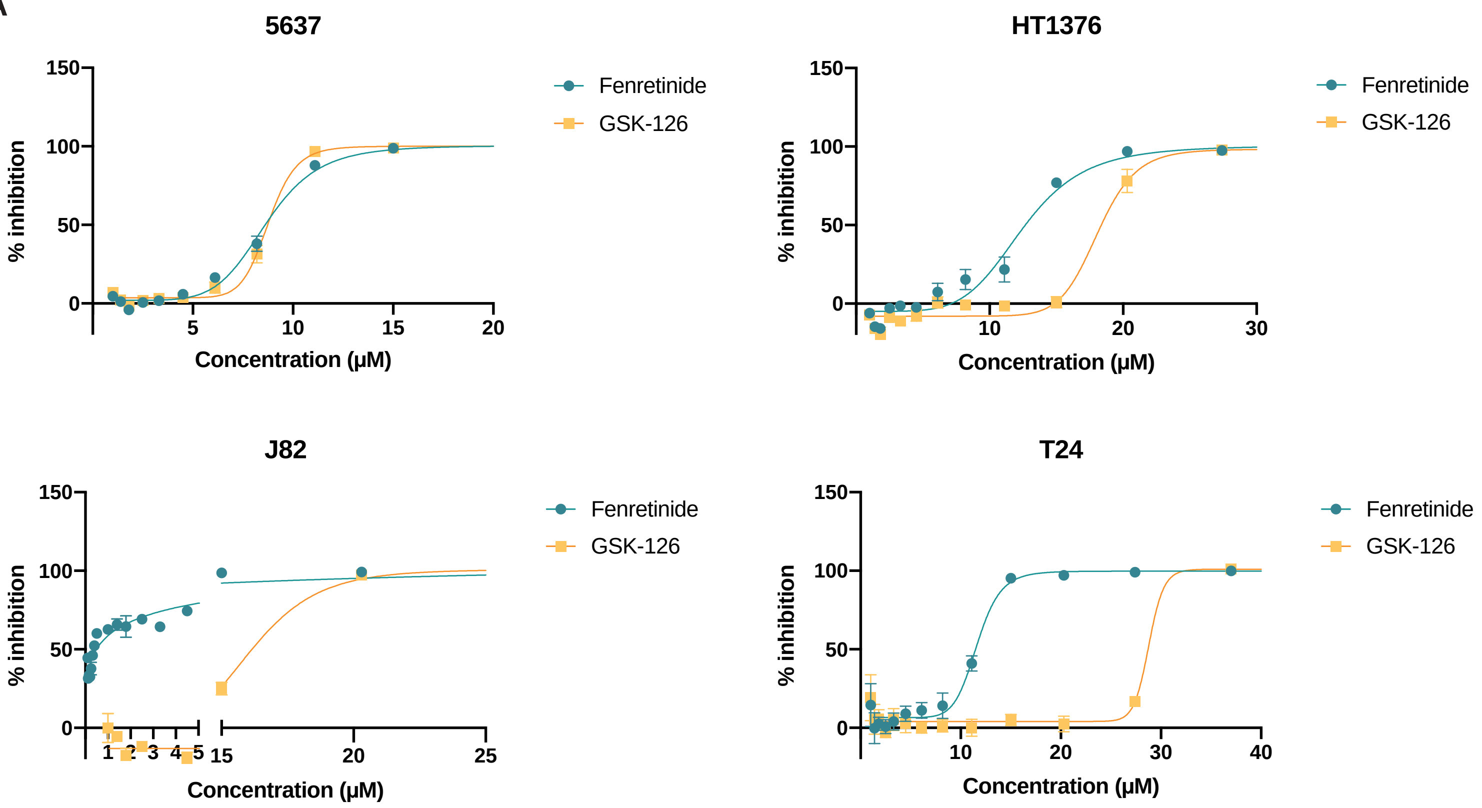


B

| IC50 (μM) | 5637 | HT1376 | J82 | T24 |
|---|---|---|---|---|
| Fenretinide | 8.67 | 12.48 | 2.17 | 11.64 |
| GSK-126 | 8.79 | 18.10 | 15.85 | 28.79 |

**Supplementary Figure 2. The effects of fenretinide and GSK-126 on MIBC cell viability**
a) Dose-dependent growth inhibition of MIBC cells upon exposure to the indicated concentrations of fenretinide or GSK-126 for 48 h. Viability was determined in comparison to untreated (DMSO only) control cells. Each point represents the mean and standard deviation of the triplicates. b) IC50 concentrations of MIBC cell lines.

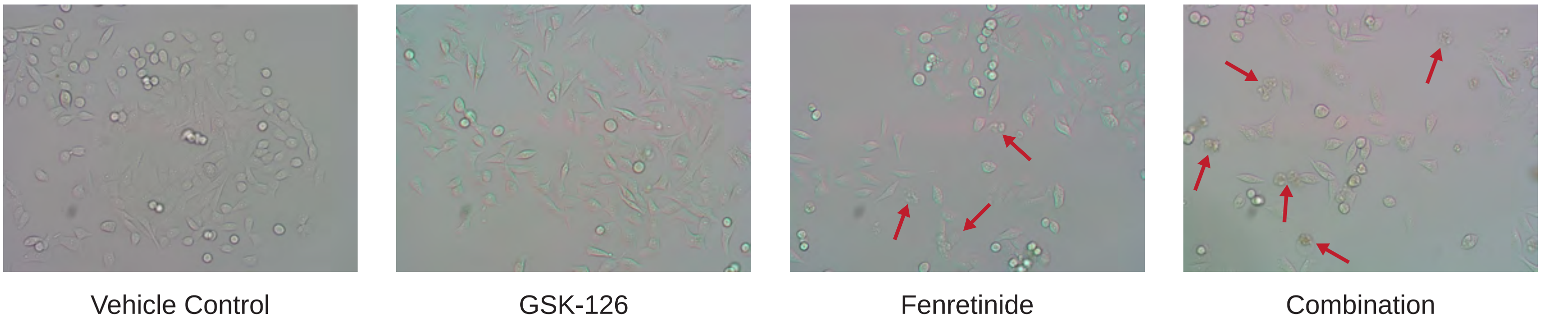


**Supplementary Figure 3. Apoptosis induced by fenretinide and GSK-126 co-treatment**
Images representing apoptosis in untreated (Vehicle control) and treated (5 μM fenretinide and 5 μM GSK-126 alone or in combination for 72h) T24 cells. Red arrows display blebs and apoptotic bodies.

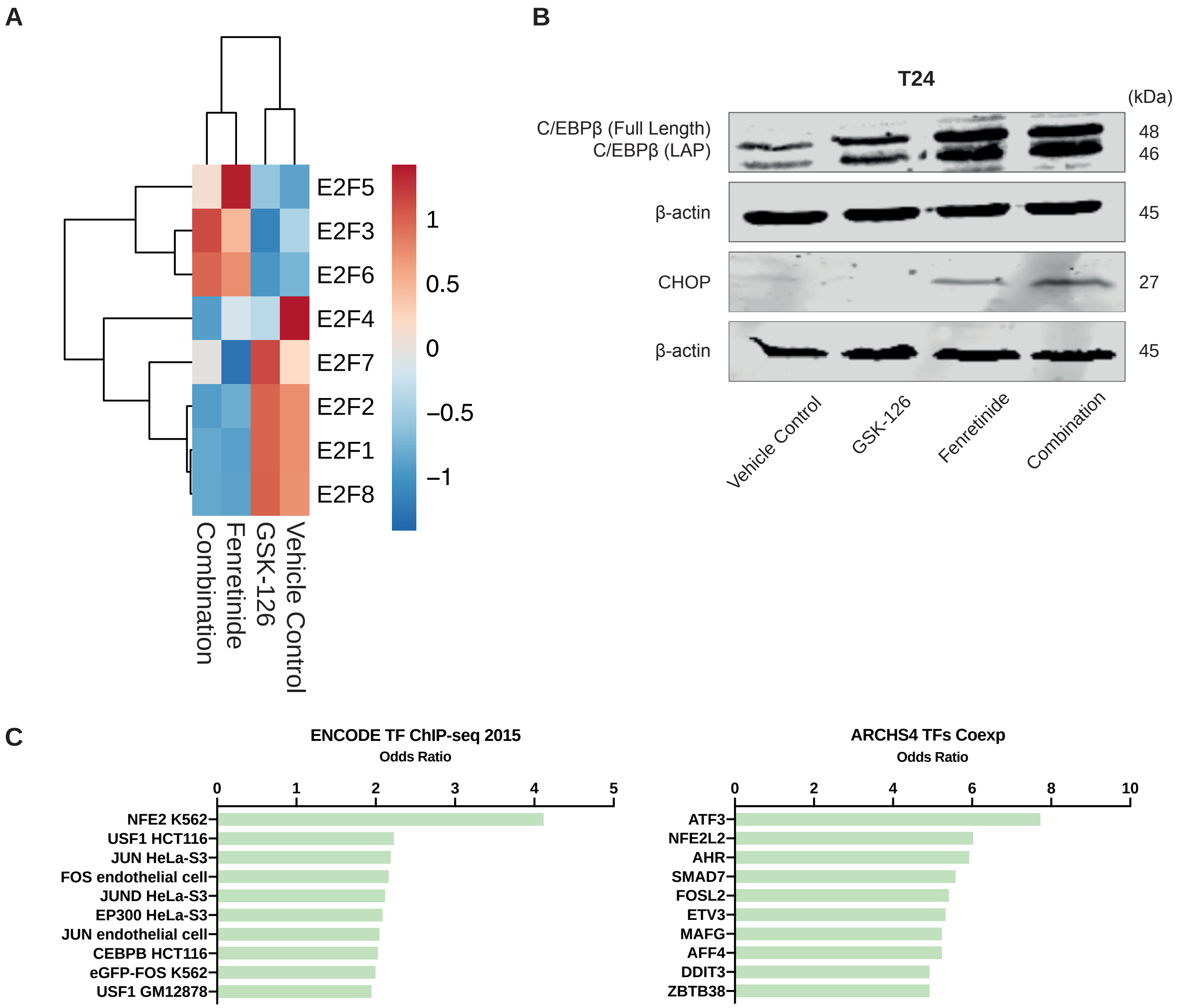


**Supplementary Figure 4. The association of fenretinide only and combination treatments with cell cycle-related processes, CEBPB and CHOP protein levels, and transcription factor activities**

a) Heat map depicting differentially expressed E2F family genes between untreated (Vehicle control) and treated (5 μM fenretinide and 5 μM GSK-126 alone or in combination for 72h) groups using fold change> 0.5 cutoff. Red and blue indicate high and low expression of genes, respectively. b) Western blots performed on T24 cell lines assessing expression of C/EBPβ and CHOP after 72 h of treatment with 5 μM fenretinide, 5 μM GSK-126, and the drug combination. c) The association of differentially expressed genes with transcription factor activities in the CoFeUR cluster was determined using the information available in the EnrichR database. Adjusted P value < 0.05.

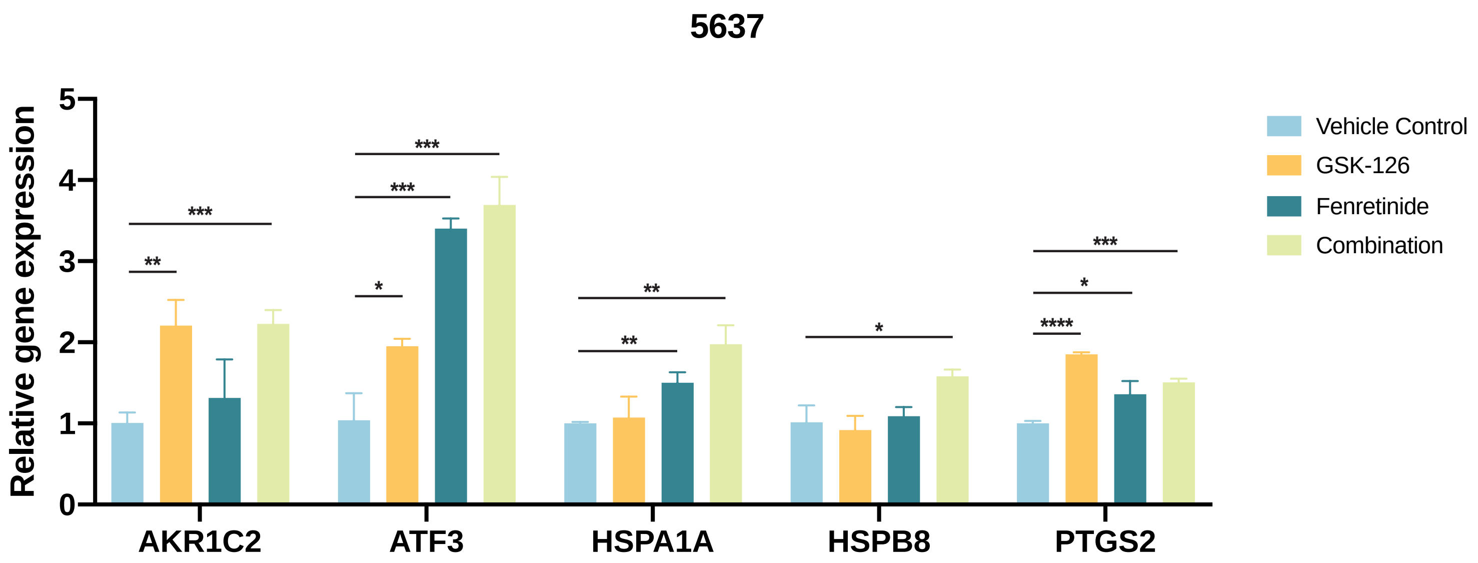


**Supplementary Figure 5. The effects of combination treatment on unfolded protein response**
The expression levels of a subset of unfolded protein response genes in 5637 cells treated with 5 μM GSK-126 and 5 μM fenretinide alone or in combination for 72h or DMSO (vehicle). Gene-specific data were normalized to GAPDH expression and are represented as average relative expression compared to vehicle controls. Error bars specify the standard deviation between triplicates. *p< 0.1, **p < 0.01, ***p < 0.001, ****p < 0.0001. p-value was determined by two-tailed, unpaired t-test.
